# Multidimensional encoding of temporal features underlies song recognition in Floridian *Ormia ochracea*

**DOI:** 10.64898/2026.05.07.723390

**Authors:** Lauren J. Bitner, Jimena A. Dominguez, Luke Bemish, Quang Vu, Julia Morgan, David A. Gray, Andrew C. Mason, Norman Lee

**Affiliations:** Department of Biology, St. Olaf College, Northfield, MN 55057, USA; Department of Biology, California State University Northridge, Northridge, CA, USA; Department of Biological Sciences, University of Toronto Scarborough, Scarborough, Ontario, M1C 1A4, Canada

**Author notes:** **Corresponding author:** Norman Lee. **Email addresses:** Lauren Bitner; Jimena Dominguez; Luke Bemish; Quang Vu; Julia Morgan; David Gray. **Senior authors:** Andrew C. Mason.

**Keywords:** temporal pattern recognition, multidimensional encoding, feature integration, acoustic communication, phonotaxis

## Abstract

Acoustic communication signals often contain complex temporal structure, yet the features underlying signal recognition remain poorly understood, particularly in eavesdropping receivers. The parasitoid fly *Ormia ochracea* localises host crickets by eavesdropping on their calling songs. In Florida, preferred host songs consist of sound pulses repeated at ~50 pulses/s, and flies exhibit matching preferences. However, it remains unclear whether this preference reflects sensitivity to individual temporal features (e.g., pulse duration, interpulse interval) or to derived temporal relationships (e.g., pulse rate, pulse period, duty cycle) that emerge from their combination. We independently varied pulse duration and interpulse interval across a broad stimulus space and quantified tethered-walking phonotaxis using a switch-following paradigm. Behavioural responses formed a structured tuning surface, with high performance along a diagonal corresponding to 50 pulses/s, as well as elevated responses for a restricted range of pulse durations across a wide range of interpulse intervals. Responses failed to collapse across stimuli sharing the same pulse rate or pulse period, indicating that these features alone do not determine recognition. Instead, behaviour was best explained by the interacting effects of pulse duration and interpulse interval. These results demonstrate that song recognition in *O. ochracea* is multidimensional, with pulse rate tuning emerging from an underlying feature space rather than a single encoded parameter.

## Introduction

Acoustic communication plays a central role in the behaviour of many animals, allowing for the transmission of information from senders to receivers (Bradbury and Vehrencamp, 1998). These signals can encode relatively static information such as species identity, sex, or individuality, as well as more dynamic information related to motivation, condition, or behavioural state (Gerhardt and Huber, 2002). Acoustic signals facilitate a wide range of behavioural interactions, including the coordination of reproductive behaviours (Andersson, 1994; Searcy and Nowicki, 2005), aggressive encounters and territorial defence (Searcy and Nowicki, 2005), predator avoidance (ter Hofstede and Ratcliffe, 2016), and prey or host localisation (Walker, 1993). For communication to function effectively in these contexts, receivers of communication signals must detect, discriminate, and recognise behaviourally relevant signals within complex natural environments (Feng and Ratnam, 2000; Miller and Bee, 2012). While these recognition processes are typically studied in the context of intended receivers (Baker et al., 2019), communication signals may also be intercepted by unintended eavesdroppers such as predators and parasitoids that exploit these signals to locate their prey or host (Bernal and Page, 2023; Zuk and Kolluru, 1998). How then do these predators and parasitoids recognise the same communication signals?

Early stages of auditory processing involve the frequency analysis of auditory input (Hedwig, 2016; Mason and Faure, 2004). However, while spectral tuning determines which signals are relevant, recognition in many systems depends critically on the extraction of the temporal structure of communication signals. Multiple conceptual frameworks have been proposed to explain temporal pattern recognition, which differ in both the computations they perform and their underlying neural implementations. One class of models emphasises feature detection, in which neurons are selectively tuned to specific temporal attributes of communication signals, such as pulse duration (Leary et al., 2008), interpulse interval (Deutsch et al., 2019; Edwards et al., 2008; Ronacher and Stumpner, 1988), or pulse rate (Schildberger, 1984; Thorson et al., 1982). Such selectivity can arise through a temporal integration process where neuronal responses accumulate over successive sound pulses, producing sensitivity to pulse rate or to a threshold number of sound pulses occurring at the appropriate interval (Alder and Rose, 1998; Rose et al., 2011). More formally, this class of computations can be described using spectrotemporal filters (Hennig, 2009), such as Gabor-like filters, which are jointly tuned to duration and temporal modulation rate and can drive temporal selectivity (Hennig et al., 2014; Ronacher et al., 2015). Recent work (Mann et al., 2025) has demonstrated that such filter-like selectivity can emerge from biologically realistic circuits through the interaction of excitation and inhibition, providing a unifying framework that links feature tuning to underlying neural dynamics. A second class of models emphasises cross-correlation or template-matching computations, in which the temporal features of signals are compared to an internal template over a defined time window (Hennig, 2003; Kostarakos and Hedwig, 2015). These models operate at the level of global pattern matching and assess the similarity between auditory input and expected temporal structure. A third class of mechanisms is based on temporal selectivity that arises from delay-line and coincidence detection mechanisms, in which inputs are temporally offset and subsequently recombined such that neurons respond maximally when signals arrive simultaneously (Jeffress, 1948; Schoneich et al., 2015). This framework provides an explicit mechanism for detecting specific interpulse intervals or pulse periods through circuit-level timing computations. While these conceptual frameworks differ in their level of description, they share the common goal of extracting temporal regularity from signals. Importantly, accumulating computational and neurophysiological evidence suggests that temporal selectivity may emerge from sequential processing involving excitatory and inhibitory interactions within local neural circuits (Clemens et al., 2021; Schoneich et al., 2015).

Despite substantial progress in understanding the neural mechanisms underlying signal recognition in intended receivers (Baker et al., 2019; Hennig et al., 2014; Kostarakos and Hedwig, 2015; Ronacher et al., 2015; Schöneich, 2020), it remains unclear whether these same mechanisms operate in unintended receivers such as predators and parasitoids that exploit communication signals. In these systems, recognition serves fundamentally different functions (locating prey or hosts vs facilitating communication between conspecifics), which may impose distinct selective pressures on sensory processing (Wikle et al., 2025). Consequently, it is unknown whether eavesdropping receivers rely on similar feature-based, interval-based, or periodicity-based mechanisms, or whether alternative strategies have evolved to extract behaviorally relevant information from communication signals.

The acoustic parasitoid fly *Ormia ochracea* is a striking example of an eavesdropper that relies on the communication signals of field crickets. Male crickets produce calling songs to attract potential mates (Alexander, 1957; Gerhardt and Huber, 2002). At the same time, gravid female flies rely on these species-specific calling songs to locate host crickets (Cade, 1975; Zuk and Kolluru, 1998) for the development of their larval young (Adamo et al., 1995; Dominguez et al., 2025). The auditory system of female *O. ochracea* is tuned to the ~5 kHz carrier frequency of host cricket calling songs, enabling detection of these songs over distances exceeding 50 meters (Robert et al., 1992; Wikle et al., 2025). Following detection, flies exhibit remarkable sound localisation abilities and can orient to sound sources with an accuracy of 2° azimuth (Mason et al., 2001). Once oriented toward a potential host, flies engage in both flying (Müller and Robert, 2001) and walking phonotaxis (Lee et al., 2009; Mason et al., 2005) to the source location. Importantly, behavioural studies demonstrate that flies can maintain preferences for host song temporal patterns across a range of ambient temperatures, consistent with physiological temperature coupling between senders and receivers (Jirik et al., 2023), although expression of this potential may be limited under some field conditions (Rossi et al., 2024). While these findings indicate that flies track temporal features of host songs under natural variation, the specific features that govern recognition remain unclear.

In Florida, *Ormia ochracea* are locally adapted to recognise and prefer the calling songs of the field cricket *Gryllus rubens* (Gray et al., 2007; Walker, 1993). These crickets produce calling songs that consist of long trills with ~10 ms pulse durations and ~10 ms interpulse intervals, resulting in a pulse rate of ~ 50 pps at an ambient temperature of 21°C (Walker, 1962; Izzo and Gray, 2004). Past experiments indicate that Floridian *O. ochracea* also exhibit preferences for a pulse rate of 50 pps (Jirik et al., 2023; Lee et al., 2019). However, it remains unclear whether this preference arises from sensitivity to individual temporal parameters, such as pulse duration or interpulse interval, or from sensitivity to derived temporal relationships, such as pulse rate, pulse period, or duty cycle that emerge from the combination of these parameters. Here, we test whether song recognition can be explained by a single temporal parameter, or whether it arises from the combined and interacting effects of multiple temporal features. By independently varying pulse duration and interpulse interval across a broad stimulus space, we assess whether behavioural responses collapse onto a single temporal axis or instead reveal multidimensional structure. More broadly, determining how eavesdropping receivers extract temporal information from communication signals provides insight into how eavesdropping receivers extract temporal information from communication signals and how these representations may differ from those of intended receivers.

## Materials and Methods

### Animals

Walking phonotaxis experiments were conducted on gravid female *Ormia ochracea* from a laboratory colony originally derived from Gainesville, Florida. The colony was maintained through manual parasitisation of *Acheta domesticus* crickets following previously described methods (Dominguez et al., 2025). Flies were reared in temperature-, humidity-, and light-controlled environmental chambers (Power Scientific Inc, model DROS52503, Pipersville, PA) set to a 12 hr light to 12 hr dark cycle at 75% humidity, and provided with butterfly nectar (The Birding Company, Yarmouth, MA, USA) *ad libitum*. A total of 52 gravid female flies were included in this study.

### Behavior Measurements

#### Experimental setup

Walking phonotaxis responses were recorded by tethering flies and placing them on top of a spherical treadmill system (Lott et al., 2007) located 25 cm away from two Avisoft Bioacoustic ultrasonic speakers (Vifa, part #60108, Germany) positioned at ± 45° azimuth to the left and right of the treadmill system relative to the forward orientation of the fly (0° azimuth). Sound presentation and treadmill data acquisition were synchronised using a National Instruments data acquisition system (NI USB-6363, USA) interfaced with a custom MATLAB (R2018a, The MathWorks Inc., USA) graphical user interface (StimProg V6; https://github.com/Ormia/Stimprog).

Locomotor data captured as x-y pixel displacements were converted to centimetres for subsequent data analyses (see below). The experimental setup was housed in an acoustically dampened sound box within a dark room kept at 30% humidity and 21 degrees Celsius.

#### Acoustic stimuli

Synthetic acoustic stimuli were generated in MATLAB using a custom script (https://github.com/Ormia/Stimprog) and then converted to analogue signals using the National Instruments data acquisition hardware (NI USB-6251, 44100 Hz), amplified (SLA1 2-Channel 140W Amplifier, USA), and broadcast through two Avisoft Bioacoustic ultrasonic speakers (Vifa, part #60108, Germany). Sound levels were controlled using programmable attenuators (Tucker Davis Technologies System 3 PA5, USA). All stimuli were calibrated at the location of the fly using a probe microphone (B&K Type 4182, Denmark) connected to a sound level meter (B&K Type 2250, Denmark). Each speaker was calibrated to 75 dB SPL (re 20 μPa). Each song consisted of a train of sound pulses; individual pulses were composed of a 5 kHz sinusoidal carrier gated by 1 ms on/off cosine-squared ramps. Pulse trains were defined by two temporal parameters: pulse duration (PD), corresponding to the total duration of each sound pulse (including the ramps), and interpulse interval (IPI), the silent interval between successive pulses. PD and IPI were independently varied across a set of values (2, 3, 5, 7, 10, 13, 15, 20, 25, 30, 35, 40, 45, and 50 ms), resulting in a total of 196 acoustic stimuli spanning a broad temporal parameter space. Sampling density varied across conditions (range: 35 to 1245 trials per stimulus), and this variation was accounted for using inverse-frequency weighting during model fitting (see below).

#### General experimental protocol

Flies were anaesthetised on ice for 5 minutes prior to experimentation and then tethered using low-melting-point wax following established methods (Jirik et al., 2023; Lee et al., 2019). Tethered flies were then mounted onto the spherical trackball system and allowed to acclimate to the behavioural testing conditions for a period of 10 minutes before testing. Experiments were conducted in a dark room under IR illumination, and behaviour was monitored using a camera equipped with a macro lens (Nikon AF MICRO NIKKOR 105 mm f/2.8 D, Japan).

Test stimuli were presented for a total of 5-seconds: 2.5-seconds from one speaker followed by a switch to the opposite speaker for 2.5 seconds (Jirik et al., 2023; Lee et al., 2019). For each individual, the direction of switching (left-to-right or right-to-left) was fixed but randomly assigned. Each experiment began with four presentations of a reference stimulus (two left-to-right and two right-to-left). The reference stimulus consisted of a synthetic song with a 10 ms PD and 10 ms IPI, modelled after the calling song of the preferred host cricket, *Gryllus rubens*, for Floridian *Ormia ochracea*. A subject was considered responsive if the total walking distance exceeded 1 cm. Only flies that responded to all initial reference stimulus presentations were included in subsequent testing. In total, 52 flies contributed to 9413 behavioural trials across the full stimulus set.

### Data analysis

The switch-following index quantified how reliably flies tracked the active sound source before and after the switch in stimulus broadcast location. Locomotor trajectories were smoothed using a moving average filter, and instantaneous angular heading was calculated from frame-to-frame changes in x-y position::

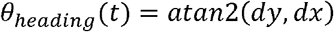

At each time point, heading alignment relative to the currently active speaker was computed as:

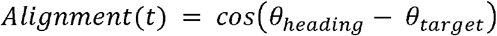

where θ target corresponded to the angular location of the active speaker (−45° before the switch and +45° after the switch, or vice versa depending on stimulus order). Alignment values were calculated separately during the pre-switch and post-switch stimulus periods, and low-velocity time points (< 0.2 cm/s) were excluded from analysis. The switch-following index was then computed as the mean of the pre-switch and post-switch alignment values, such that higher scores indicate stronger tracking of the active speaker across the speaker transition.

The switch-following index was then calculated as:

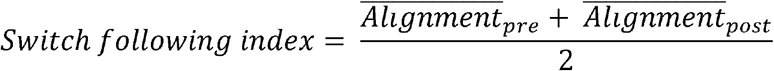

where 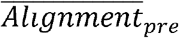 and 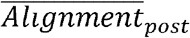 represent the mean alignment values during the pre-switch and post-switch stimulus periods, respectively. Higher switch-following index values indicate stronger tracking of the active speaker across the speaker transition.

To test alternative hypotheses about song recognition, we modelled switch-following performance as a function of pulse duration and interpulse interval using generalised additive mixed models (GAMMs; mgcv package in R). Pulse duration and interpulse interval were treated as continuous variables, allowing behavioural responses to be estimated as a smooth response surface across the stimulus space. Individual trials were included in the analysis, with fly identity included as a random intercept to account for repeated measurements

As sampling density varied across the PD and IPI stimulus grid (some conditions more frequently tested than others), we applied inverse-frequency weights such that each PD and IPI combination contributed approximately equally to model estimation. All candidate models were fit using identical weighting to ensure valid model comparison.

We compared a set of candidate models representing alternative hypotheses about the temporal features underlying song recognition. These included: (1) a joint feature-integration model incorporating a two-dimensional tensor product smooth of PD and IPI, (2) an additive model including independent smooth terms for PD and IPI, (3) a pulse-period model including a smooth function of pulse period (PD + IPI) with an additional PD term, (4) a pulse-period-only model, and (5) models based on duty cycle (PD / [PD + IPI]), including a duty-cycle-only model and models combining duty cycle with pulse period, with and without an interaction term. Models were fitted using maximum likelihood (ML) for comparison and ranked using Akaike’s Information Criterion (AIC). The best-supported model was then refitted using restricted maximum likelihood (REML) for parameter estimation and visualisation. To assess robustness, analyses were repeated after excluding the oversampled condition (PD = 10 ms, IPI = 10 ms).

All behavioural data preprocessing and initial analyses were performed in MATLAB (R2019a, The MathWorks Inc., USA). Statistical analyses and model fitting were conducted in R (version 4.5.3).

## Results

### Switch-following index captures song preference behaviour

To validate the switch-following index, we compared phonotactic responses in response to the preferred 50 pulses per second song and the nonpreferred 10 pps song (Lee et al., 2019). Fly-level mean responses were calculated for each stimulus condition (n = 52 flies for 50 pps; n = 34 for 10 pps). In response to the 50 pps stimulus, flies exhibited consistent phonotactic tracking of the active speaker, aligning their angular heading with the pre-switch target location and rapidly reorienting following the change in broadcast location (Fig 1A). In contrast, responses to the 10 pps stimulus were more variable and lacked clear tracking of the broadcasting speakers (Fig 1B). These differences in behavioural dynamics were captured by the switch-following index, which was significantly higher for the 50 pps stimulus compared to the 10 pps stimulus (Fig. 1C; Welch’s t-test, t(65.94) = 7.066, p = 1.241e-09). Therefore, these results demonstrate that the switch-following index reliably quantifies stimulus-dependent differences in phonotactic tracking and provides a robust measure of song recognition.

**Figure 1.**
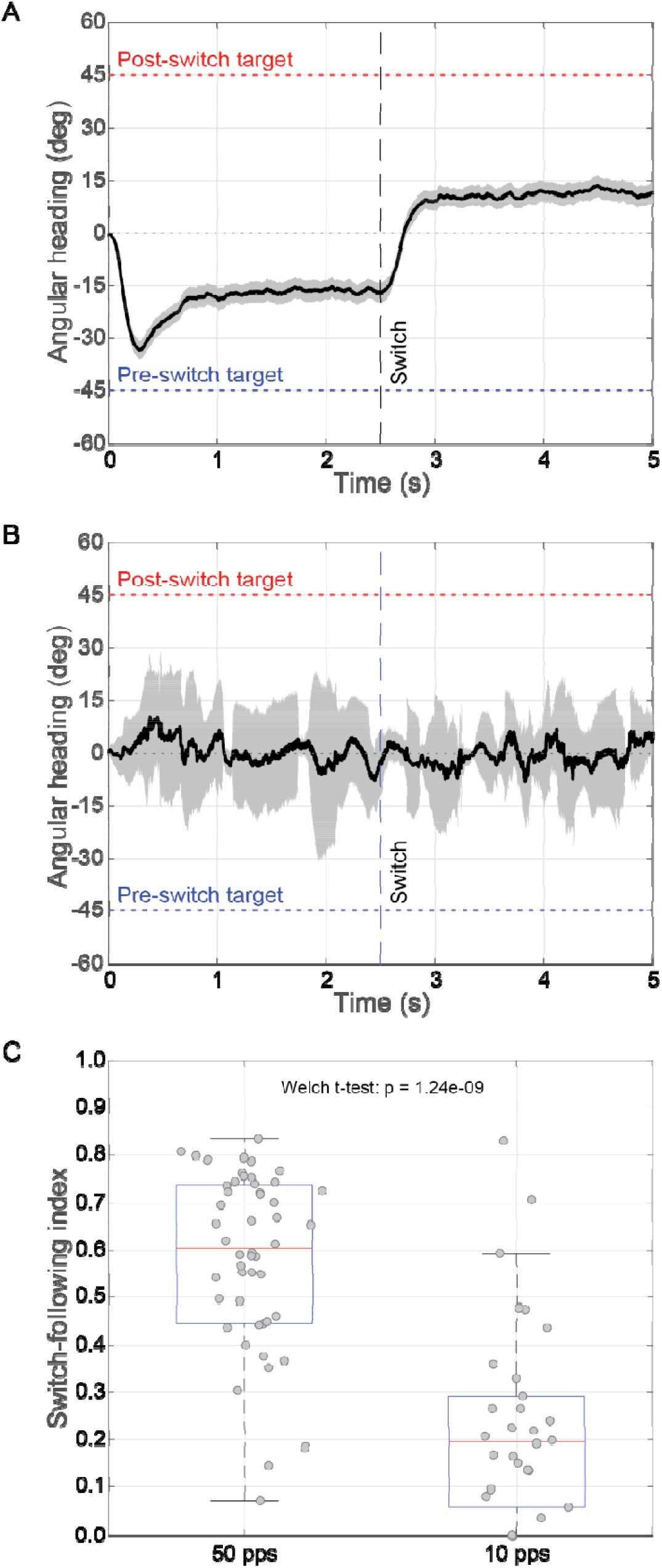
Switch-following paradigm reveals song-specific tracking behaviour. (A) Mean angular heading of flies in response to a preferred 50 pulses per second (pps) stimulus. Flies initially orient toward the pre-switch target (−45°, blue dashed line) and rapidly reorient toward the post-switch target (+45°, red dashed line) following the speaker switch (vertical dashed line at 2.5 s). Black trace shows the across-fly mean angular heading; grey shading indicates 95% CI. (B) Mean angular heading in response to a non-preferred 10 pps stimulus. Flies show weak and inconsistent tracking of the active speaker, with little evidence of reorientation following the switch. (C) Switch-following index (SFI) quantified for individual flies under each stimulus condition. Points represent fly-level mean SFI values; boxplots show median and interquartile range. Flies exhibited significantly higher SFI values in response to the 50 pps stimulus compared to the 10 pps stimulus, indicating stronger tracking of the active speaker for behaviorally relevant song patterns.

### Behavioural responses form a structured tuning surface in pulse duration interpulse interval space

To visualise how different combinations of PD and IPI jointly shape song recognition, we constructed a two-dimensional heatmap of switch-following index scores across the full range of PD and IPI combinations (Fig. 2A). Heatmap values represent across-fly means (n = 52 flies; number of trials per condition = 35 to 1245). Behavioural responses were not uniformly distributed across the stimulus space, but instead exhibited a structured region of high performance. Flies exhibited the strongest switch-following for PDs of ~5–15 ms paired with IPIs of ~15–35 ms.

**Figure 2.**
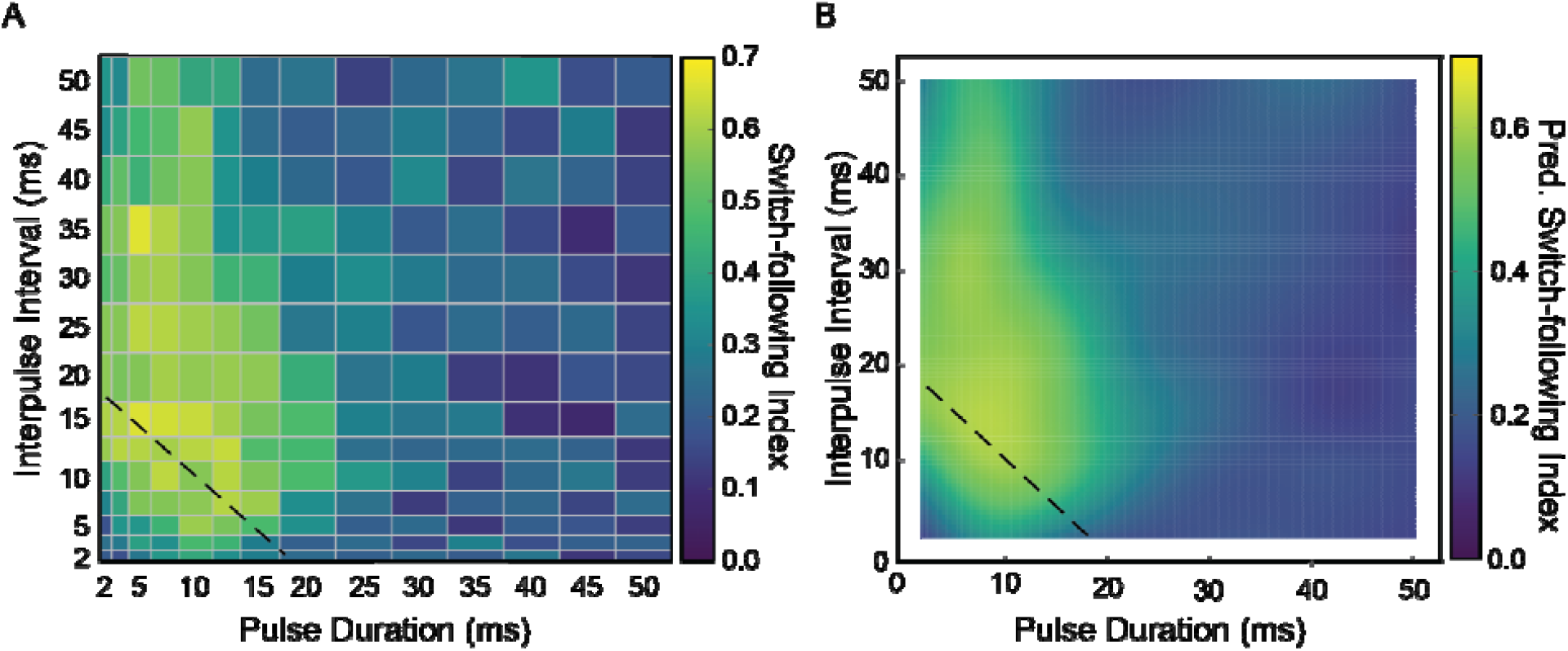
Behavioural tuning reveals multidimensional encoding of temporal features. (A) Heatmap displays across-fly mean switch-following score as a function of pulse duration (PD, x-axis) and interpulse interval (IPI, y-axis). Warmer colours indicate stronger phonotactic tracking of the active speaker following a switch in the broadcast location. Responses form a structured tuning surface with peak performance near PD-IPI combinations corresponding to ~ 50 pulse/sec (dashed diagonal line), as well as elevated responses across a restricted range of PD spanning a range of IPI. (B) Heatmap displays model-predicted switch-following scores from the best-supported generalised additive mixed model (GAMM), including a smooth interaction between pulse duration and interpulse interval. The fitted surface recapitulates the structure observed in the empirical data, including the diagonal band of high performance values and the extended region of elevated responses across a restricted range of pulse durations. These results indicate that behavioural responses depend on combinations of temporal features rather than a single parameter.

Notably, elevated performance scores were distributed along a diagonal band of PD-IPI combinations that preserve a similar pulse rate (Fig. 2A dashed diagonal, ~50 pps), giving the appearance of tuning to pulse rate. However, responses were not strictly constrained to this diagonal. Intermediate performance was observed across a restricted range of PD (~5–10 ms) spanning a broad range of IPI, indicating sensitivity to PD independent of pulse period.

Together, these patterns suggest that while behavioural responses exhibit structure consistent with pulse period tuning, they are not fully explained by pulse period tuning alone. Instead, these response patterns may reflect the combined influences of PD and IPI.

### Model comparison supports joint encoding of temporal features

Model comparison strongly supported models incorporating interactions between temporal features over models based on single parameters (Table 1). The tensor-product smooth model, including PD and IPI, provided an excellent fit to the data (AIC = 6436.0). A model based on the interaction between duty cycle and pulse period also performed similarly well (ΔAIC < 1), indicating that these alternative parameterisations capture similar structure in the behavioural responses.

**Table 1.**
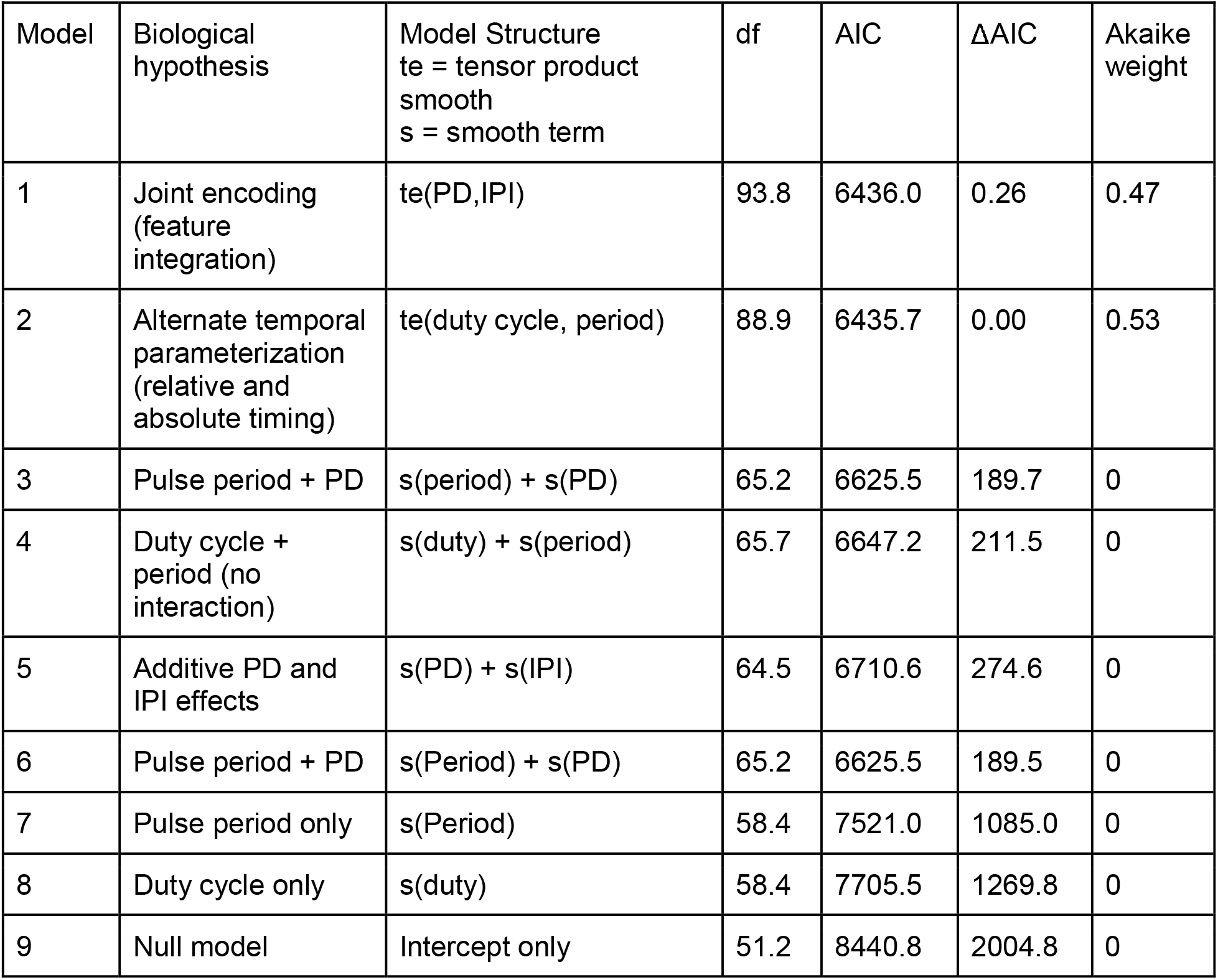
Comparison of candidate models representing alternative hypotheses for song temporal feature coding. Comparison of candidate models representing alternative hypotheses for song temporal feature coding. Models incorporating interactions were fit as generalized additive mixed models with a random intercept for fly identity. Pulse duration (PD) and interpulse interval (IPI) were treated as continuous predictors. Inverse-frequency weights were applied to account for unequal sampling density across the stimulus grid. Model fit was evaluated using Akaike’s Information Criterion (AIC), with ΔAIC calculated relative to the best-supported model. Models incorporating interactions between temporal features received the strongest support. A joint PD–IPI response-surface model and a model parameterized using duty cycle and pulse period performed comparably (ΔAIC < 1), indicating that the behavioural response surface can be similarly described using either temporal parameterization.

| Model | Biological hypothesis | Model Structure<br>te = tensor product<br>smooth<br>s = smooth term | df | AIC | $\Delta\text{AIC}$ | Akaike weight |
| --- | --- | --- | --- | --- | --- | --- |
| 1 | Joint encoding (feature integration) | te(PD,IPI) | 93.8 | 6436.0 | 0.26 | 0.47 |
| 2 | Alternate temporal parameterization (relative and absolute timing) | te(duty cycle, period) | 88.9 | 6435.7 | 0.00 | 0.53 |
| 3 | Pulse period + PD | s(period) + s(PD) | 65.2 | 6625.5 | 189.7 | 0 |
| 4 | Duty cycle + period (no interaction) | s(duty) + s(period) | 65.7 | 6647.2 | 211.5 | 0 |
| 5 | Additive PD and IPI effects | s(PD) + s(IPI) | 64.5 | 6710.6 | 274.6 | 0 |
| 6 | Pulse period + PD | s(Period) + s(PD) | 65.2 | 6625.5 | 189.5 | 0 |
| 7 | Pulse period only | s(Period) | 58.4 | 7521.0 | 1085.0 | 0 |
| 8 | Duty cycle only | s(duty) | 58.4 | 7705.5 | 1269.8 | 0 |
| 9 | Null model | Intercept only | 51.2 | 8440.8 | 2004.8 | 0 |

Consistent with the empirical tuning surface, the fitted PD–IPI response surface closely recapitulated the structure observed in the empirical heatmap, including the diagonal ridge and extended region of elevated responses that are not aligned with either PD or IPI axes (Fig 2B), indicating that this structure arises from the interaction between temporal features rather than from independent effects of either parameter. The model explained a substantial proportion of variance in behavioural responses (adjusted R^2^ = 0.33), and the smooth interaction between PD and IPI was highly significant (p < 0.001). Trial order had no significant effect on behaviour (p = 0.56).

In contrast, models based on single temporal parameters performed substantially worse. A pulse-period-only model showed poor fit (ΔAIC = 1085.0), and a duty-cycle-only model also performed poorly, suggesting that relative pulse structure alone is insufficient to account for behaviour. Adding PD to the pulse-period model improved fit (ΔAIC = 189.5), but this model still performed relatively poorly compared to models that included interactions between temporal features. Likewise, an additive model including independent effects of PD and IPI was insufficient (ΔAIC = 274.6), indicating that the interaction between temporal features is critical for explaining behaviour.

Together, these results indicate that song recognition is best explained by the combined and interacting effects of temporal features rather than by any single parameter. Although behavioural responses could be similarly described using either PD/IPI or duty cycle/pulse period parameterisations, both models required interactions between temporal features, consistent with a multidimensional feature integration mechanism.

### Pulse duration and interpulse interval effects are interdependent

Consistent with the model results, we next examined behavioural responses along each temporal dimension while holding the other constant (Fig. 3).

**Figure 3.**
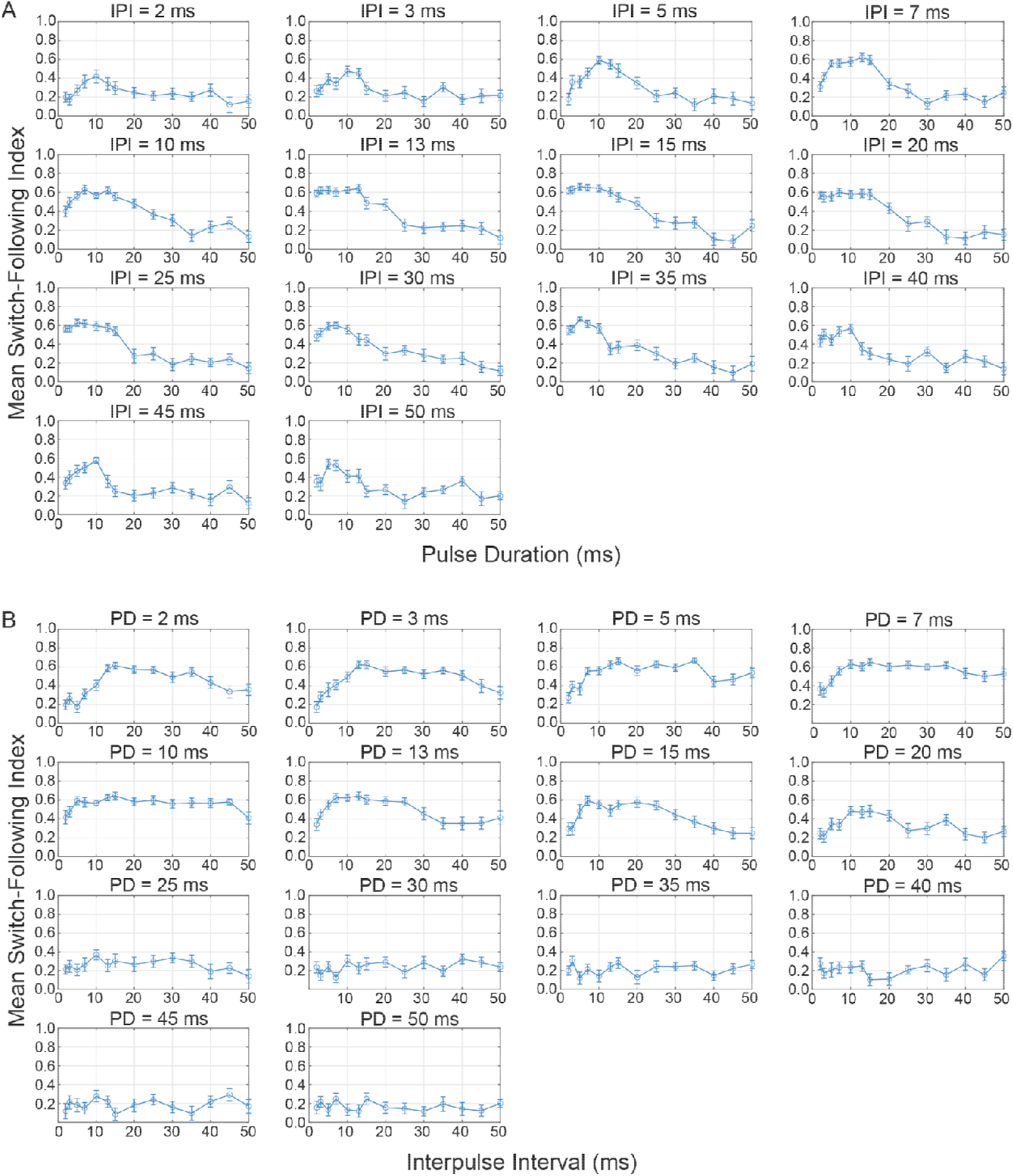
Pulse duration and interpulse interval tuning are interdependent. (A) Mean switch-following scores plotted as a function of pulse duration (PD) for each interpulse interval (IPI). Each panel shows responses at a fixed IPI (2-50 ms), with PD varying along the x-axis. Points represent across-fly means (n = 52 flies), and error bars indicate ± SEM across flies. PD tuning varies systematically with IPI, indicating that behavioural responses depend on combinations of temporal features rather than a single parameter. (b) Mean switch-following scores plotted as a function of interpulse interval (IPI) for each interpulse interval (IPI). Each panel shows responses at a fixed PD (2-50 ms), with IPI varying along the x-axis. Points represent across-fly means (n = 52 flies), and error bars indicate ± SEM across flies. IPI tuning varies systematically with PD, with elevated responses extending across a broad range of intervals, indicating multidimensional sensitivity to temporal features.

To assess sensitivity to PD, we examined behavioural responses as a function of PD while holding IPI constant (Fig. 3A). At short to intermediate IPIs (~5–20 ms), flies exhibited clear tuning to PD with peak switch-following performance at PD of ~5–15 ms. However, this tuning was not consistent across all IPI. As IPI increased, overall response magnitudes declined and tuning curves flattened out, with little evidence of PD selectivity at longer IPIs. These results indicate that PD tuning is contingent on IPI and cannot fully account for song recognition behaviour.

We next examined behavioural responses as a function of IPI while holding PD constant (Fig. 3B). For intermediate PDs (~5–15 ms), flies responded robustly across a broad range of interpulse intervals (~10–30 ms), resulting in relatively flat tuning profiles. In contrast, for longer PDs (>20 ms), responses were uniformly low across all IPIs. Similarly, very short PDs elicited weak and inconsistent responses. These results suggest that the IPI alone does not determine song recognition.

### Responses do not collapse across iso rate and iso-period stimulus combinations

To directly test whether song recognition can be explained by a single temporal parameter, we examined responses across iso-rate and iso-period stimulus combinations (Fig. 4). Behavioural responses across stimuli with the same pulse rate (Fig 4A) or pulse period (Fig 4B), which contradicts predictions of a single parameter encoding model. Although mean responses exhibited peaks near 50 pps Hz and ~20 ms, respectively, substantial variability was observed among different combinations of PD and IPI that produced the same rate or period. In particular, responses at ~50 pps varied widely across stimulus combinations.

**Figure 4.**
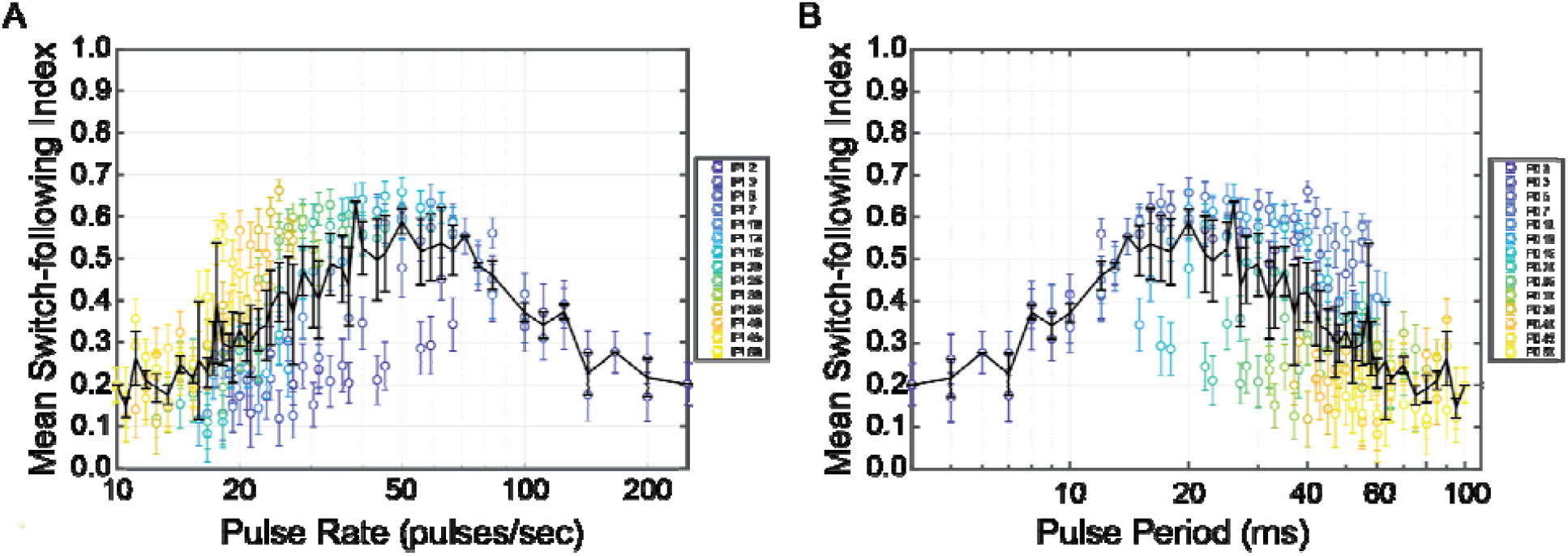
Tests of iso-rate and iso-period collapse. (A) Mean switch-following score plotted as a function of pulse rate (Hz). Each point represents a unique combination of pulse duration (PD) and interpulse interval (IPI) averaged across flies (n = 52 flies) that yields the same pulse rate, with colours indicating different pulse durations. The black line shows the mean response across all combinations at each pulse rate (± SEM). (B) Mean switch-following score plotted as a function of pulse period (ms; PD + IPI). As in (A), points represent different PD–IPI combinations that produce the same pulse period, with colours indicating pulse duration. The black line indicates the mean response (± SEM) across all combinations at each pulse period. Responses do not converge across iso-rate or iso-period stimuli, indicating that behavioural responses depend on specific combinations of pulse duration and interpulse interval rather than a single temporal parameter.

These results highlight a key limitation of one-dimensional representations of the stimulus space. Plotting responses as a function of pulse rate or pulse period collapses across distinct PD-IPI combinations, thereby obscuring variability that reflects sensitivity to specific combinations of temporal features. As a result, these representations can give the appearance of tuning to a single parameter while masking the underlying multidimensional structure of behavioural responses. Together, these findings demonstrate that song recognition cannot be reduced to pulse rate, pulse period, or other derived one-dimensional features, but instead depends on the combined and interacting effects of PD and IPI.

## Discussion

Our results demonstrate that temporal pattern recognition in *Ormia ochracea* cannot be explained by a simple pulse rate detection mechanism. Consistent with prior work (Jirik et al., 2023; Lee et al., 2019; Walker, 1993), phonotactic performance peaked near 50 pps (and for a corresponding pulse period of 20 ms), but failed to collapse along different pulse duration and inter-pulse interval combinations resulting in the same preferred pulse rate (Fig 4). This lack of iso-pulse rate collapse indicates that stimuli with identical pulse rates or pulse periods are not functionally equivalent to the auditory system, which argues against song recognition models based purely on pulse periods or autocorrelation-like computations. In the Floridian *Ormia ochracea*, song temporal pattern recognition cannot be reduced to the acoustic feature extraction of a single scalar variable such as pulse rate or pulse period.

Instead, multiple lines of evidence support a multidimensional encoding of temporal features. First, both the empirical heatmap and the model-predicted response surface (Fig 2), together with conditional tuning analyses (Fig. 3), reveal structured tuning that cannot be captured by a single axis. Importantly, the fitted surface reproduces features that are not aligned with either temporal dimension, and thus these patterns arise from nonlinear interactions between pulse duration and interpulse interval rather than independent tuning to either feature. While there is a diagonal band of relatively high response scores describing a preferred range of pulse rates or pulse periods, we also see clear deviations from this diagonal. Second, responses do not collapse across stimuli that share the same pulse rate or pulse period (Fig 4), indicating that these derived temporal relationships alone are insufficient to explain recognition. Third, our statistical models demonstrate that including pulse duration, interpulse interval, and their interaction provides a significantly better fit to the data than the alternative model based solely on pulse period (Table 1). Notably, the significance of the pulse duration by inter-pulse interval interaction term suggests that these features combine nonlinearly in modulating behaviour. Together, these results show that the auditory system retains sensitivity to the individual temporal components of the host cricket calling song (pulse durations and inter-pulse intervals), and integrates these features for pulse rate sensitivity to guide phonotactic behaviour.

These results constrain the underlying neural computation. Classical models of temporal processing, such as delay-line and coincidence detection circuits (Jeffress, 1948), provide a useful starting point for understanding the neural mechanisms of song pattern recognition (Kostarakos and Hedwig, 2015; Schoneich et al., 2015). In terms of pulse rate recognition, an acoustic feature detector sensitive to pulse rates may arise from detectors that respond best when subsequent sound pulses arrive simultaneously at a coincidence detector. This can be achieved if auditory neurons physiologically introduce a temporal delay matching the salient interpulse interval for a specific preferred pulse rate (Schoneich et al., 2015). However, such a model for pulse rate recognition would predict strong invariance across stimuli sharing the same pulse rate, which is inconsistent with our data (Fig 4). More generally, computational modelling of song recognition circuits in insects has demonstrated that tuning to specific temporal features, including pulse duration, interpulse interval, or pulse rate, can emerge from modifications of circuit parameters even in the absence of an explicit coincidence detection mechanism (Clemens et al., 2021). This suggests that similar tuning functions do not uniquely specify the underlying neural mechanism. One possibility is that temporal pattern processing may occur across multiple stages (Hennig et al., 2014; Schildberger, 1984), in which early stages involve the encoding of pulse duration and interpulse interval separately, and later stages of processing integrate these features through the interplay of excitation and inhibition (Edwards et al., 2008; Leary et al., 2008) or temporally offset integration windows (Sarmiento-Ponce et al., 2018; Schoneich et al., 2015). Such architectures could give rise to the observed non-linear interactions between pulse duration and interpulse interval while maintaining sensitivity to these underlying temporal features rather than reducing them to a single derived metric such as pulse rate or pulse period.

While song pulse rate influences the phonotactic behaviour of Floridian *Ormia ochracea*, failure of iso-rate and iso-period collapse indicates that pulse rate tuning may not be explicitly encoded by the auditory nervous system. Instead, we propose that pulse rate preferences emerge as a projection of the underlying multidimensional feature space. Specifically, different combinations of pulse duration and interpulse interval that fall within a behaviorally permissive region of this response space will necessarily correspond to a range of pulse rates and apparent pulse rate preference when phonotactic responses are plotted along this single dimension (Jirik et al., 2023; Lee et al., 2019; Walker, 1993). In this sense, pulse rate can be a useful descriptive variable, but it does not capture the full structure of the computation. This interpretation highlights the limitations of one-dimensional preference functions (Kilmer et al., 2017), which can obscure preferences based on multivariate integration, or any latent preferences not revealed by a single feature (Gray et al., 2016).

Temporal pattern recognition has been studied in the intended receivers of a range of acoustically communicating species (Baker et al., 2019; Gerhardt and Huber, 2002). In many of these systems, signal recognition is described in terms of sensitivity to specific temporal parameters such as pulse duration (Hennig, 2003), interpulse interval, and pulse rate (Doherty, 1985; Hennig, 2003; Thorson et al., 1982). For example, in two closely related cricket species, *Teleogryllus oceanicus* are selective for a range of pulse rates, while *Teleogryllus commodus* exhibit selectivity for pulse duration (Hennig, 2003). Similarly, female Cope’s grey treefrog *Hyla chrysoscelis* also exhibits selectivity for advertisement calls with pulse rates that match those produced by conspecific males (Lee and Mason, 2017; Schul and Bush, 2002). However, not all intended receivers can be described by tuning to a single temporal parameter. In some species, behavioural responses span a broad range of pulse duration and interpulse interval combinations, which suggests sensitivity to a combination of temporal features rather than a single derived metric. For instance, despite male *Hyla versicolor* producing advertisement calls with a specific pulse rate, females exhibit preferences for a broad range of pulse duration and interpulse intervals that comprise a broad rectangular recognition space (Schul and Bush, 2002). Similarly, in Floridian *Ormia ochracea*, rather than observing a narrow diagonal band of performance scores around a particular pulse rate preference, deviations away from this band indicate sensitivity to a combination of values. Thus, temporal pattern recognition can range from relatively low-dimensional parameter tuning to more complex multi-dimensional integration of temporal features.

Several limitations of the present study point to future directions. First, since our conclusions are based on behavioural data, direct tests of mechanisms of song recognition will depend on neurophysiological recordings combined with neuroanatomical results throughout the auditory pathway. Both female *O. chracea* and female crickets depend on recognising and localising the same cricket calling songs to find their host or mate, respectively. Taking a comparative approach to characterise similarities or differences in their auditory processing can help shed light on common or diverse solutions to shared problems (Gerhardt and Huber, 2002). Second, *O. ochracea* occur across parts of the southern US, the Hawaiian Islands, and in Mexico (Gray et al., 2019; Lehmann, 2003). As *O. ochracea* from different regions are locally adapted to prefer different host cricket species singing songs with drastically different temporal features (Gray et al., 2007), it remains unknown whether song recognition is based on the same combination of temporal features described in the current study. Third, the host cricket calling songs of *T. oceancius* have been found to be rapidly evolving in response to selection from flies (Gallagher et al., 2023; Tinghitella et al., 2018; Tinghitella et al., 2021). These novel songs are more cryptic and differ in the frequency content and fine-scale temporal features compared to the ancestral calling songs (Tinghitella et al., 2018). Likely in response to host availability or the cryptic nature of novel songs, some populations of Hawaiian *O. ochracea* have also been found to switch hosts and to rely on multiple host species (Broder et al., 2023). These changes suggest that some level of flexibility in recognising temporal features would be adaptive for Hawaiian *O. ochracea*.

In summary, our results demonstrate that temporal pattern recognition in Floridian *Ormia ochracea* cannot be explained by a simple pulse rate detection mechanism. Instead, phonotactic behaviour indicates sensitivity to multiple temporal features, which include: pulse duration, interpulse interval, and their non-linear interaction. This multidimensional structure constrains the underlying neural computation such that it is unlikely to be a single derived variable, but emerges from the integration of distinct temporal components. Within this framework, apparent pulse rate tuning arises as a projection of an underlying feature space rather than an explicitly encoded attribute. More broadly, our findings highlight the importance of considering multidimensional representations when studying signal recognition. Similar principles may underlie pattern recognition across a range of acoustic communication systems, including both intended receivers and eavesdroppers.

## Acknowledgements

We would like to thank The Bug Company (Ham Lake, MN), especially their staff, for providing us with a constant, reliable source of *Acheta domesticus* that supports research activities in the Lee Lab at St. Olaf College. We thank past and present members of Lee Lab for help with animal care and propagating fly colonies. Funding was provided by a National Science Foundation CAREER grant to NL (IOS 2144831), with additional support from the St. Olaf Collaborative Undergraduate Research and Inquiry (CURI) Program

## Author contributions according to the <u>CRediT system</u>

LJB: conceptualisation (supporting), formal analysis (supporting), investigation (lead), methodology (supporting), Visualization (equal), writing – original draft (equal), and writing – review & editing (equal)

J.A.D.: investigation (equal) and writing – review & editing (supporting).

LB: investigation (supporting), formal analysis (supporting), visualisation (supporting), writing – review & editing (equal).

Q.V.: investigation (supporting) and writing – review & editing (supporting). J.M.: investigation (supporting)

D.A.G: conceptualisation (supporting), methodology (supporting), writing – review & editing (equal)

A.C.M: conceptualisation (supporting), formal analysis (supporting), funding acquisition (supporting), investigation (supporting), methodology (equal), project administration (equal), writing – review & editing (supporting).

N.L: conceptualisation (lead), data curation (lead), formal analysis (equal), funding acquisition (equal), investigation (supporting), methodology (equal), project administration (equal), Resources (lead), software (lead), Supervision (lead), Validation (lead), Visualization (lead), writing – original draft (equal), and writing – review & editing (equal).

